# TAPIOCA: Topological Attention and Predictive Inference of Chromatin Arrangement Using Epigenetic Features

**DOI:** 10.1101/2021.05.16.444378

**Authors:** Max Highsmith, Jianlin Cheng

**Affiliations:** Department of Electrical Engineering and Computer Science, University of Missouri, Columbia, MO 65211, USA

## Abstract

Chromatin conformation is an important characteristic of the genome which has been repeatedly demonstrated to play vital roles in many biological processes. Chromatin can be characterized by the presence or absence of structural motifs called topologically associated domains. The de facto strategy for determination of topologically associated domains within a cell line is the use of Hi-C sequencing data. However Hi-C sequencing data can be expensive or otherwise unavailable. Various epigenetic features have been hypothesized to contribute to the determination of chromatin conformation. Here we present TAPIOCA, a self-attention based deep learning transformer algorithm for the prediction of chromatin topology which circumvents the need for labeled Hi-C data and makes effective predictions of chromatin conformation organization using only epigenetic features. TAPIOCA outperforms prior art in established metrics of TAD prediction, while generalizing across cell lines beyond those used in training.

**Availability:** the source code of TAPIOCA and training and test datasets are available at https://github.com/Max-Highsmith/TAPIOCA

**Author Summary:** In this paper we outline a machine learning approach for predicting the topological organization of chromosomes using epigenetic track data as features. By utilizing an architecture inspired by the sequence transduction transformer network we are able to effectively predict multiple metrics used to characterize topologically associated domains. Our experimental results demonstrate that once trained our algorithm can effectively predict topological organization on novel cell lines all without any exposure to original Hi-C data in test datasets.

## Introduction

As the cost of genetic sequencing continues to decrease at a rate surpassing Moore’s law (“DNA Sequencing Costs: Data” n.d.), the plethora of available sequencing data has permitted the wide-spread application of machine learning approaches to the field of genomics. These approaches have spanned a variety of applications including: Assay denoising (Hong et al. 2020) (Dimmick, Lee, and Frey 2020) (Highsmith and Cheng 2020). 3D modeling (Oluwadare, Highsmith, and Cheng 2019), and regulatory network prediction (Razaghi-Moghadam and Nikoloski 2020). In this paper we outline the application of a recently developed machine learning algorithm, the Transformer, to the task of topologically associated domain (TAD) identification using epigenetic features as a proxy for Hi-C data.

The three dimensional genome has repeatedly been revealed to play an important role in a plethora of important biological processes. One prolific assay for inspecting the 3D organization of the genome is Hi-C, a variant of chromosome conformation capture (3C) assay. Hi-C data can, in addition to many other applications, be used to identify regions of the genome with preferentially self-interacting regions termed topologically associated domains (TADs). Substantial scientific attention has been directed at the development of tools for identifying TAD’s using Hi-C data (Zufferey et al. 2018) (Dixon et al. 2012) (Filippova et al. 2014).

It has been demonstrated that epigenetic features such as repressive histone modifications have preferential association with inter-TAD boundaries, indicating the potential contribution of epigenetic modifications in the construction of TADs. TAD prediction using epigenetic features was first formulated as a classification problem using logistic regression to predict boundaries(Ulianov et al. 2016). The approach was later expanded to include lasso regression and gradient boosting (Ramírez et al. 2018). The task was then reformulated as a linear regression problem in which epigenetic features were used to predict transitional gamma, a continuous metric created by the authors based on TAD identification tool Armatus. Recently the first application of neural networks, specifically LSTM obtained state-of-the are predictions (Rozenwald et al. 2020)

This paper provides two important contributions relative to prior research in this area. First we build upon the use of machine learning methods by applying self-attention through a variant of the state-of-the-art Transformer model which we call TAPIOCA (**T**opological **A**ttention and **P**redictive **I**nference of **C**hromatin **A**rrangement). Second we extend the metrics for TAD characterization beyond the previously used transitional gamma to incorporate more prolific metrics for TAD characterization such as Insulation Score (Crane et al. 2015) and Directionality Index (Dixon et al. 2012). Through these extensions and comparative analysis to the results of previously suggested models, we strengthen the case for dependence between epigenetic profile and TAD formation.

## Results

### Overview of Dataset Features and Labels

Previous work on prediction of topological organization in Drosophila based on epigenetic features has used a metric called Transitional gamma. Transitional gamma is computed by performing TAD calling using the armatus tool with gamma values 1-10 and assigning the transitional gamma of a loci to be the first gamma value at which armatus identifies a TAD boundary (Figure 1a). In our experiments we use the transitional gamma values assigned to the feature dataset by (Rozenwald et al. 2020).

**Figure 1.**
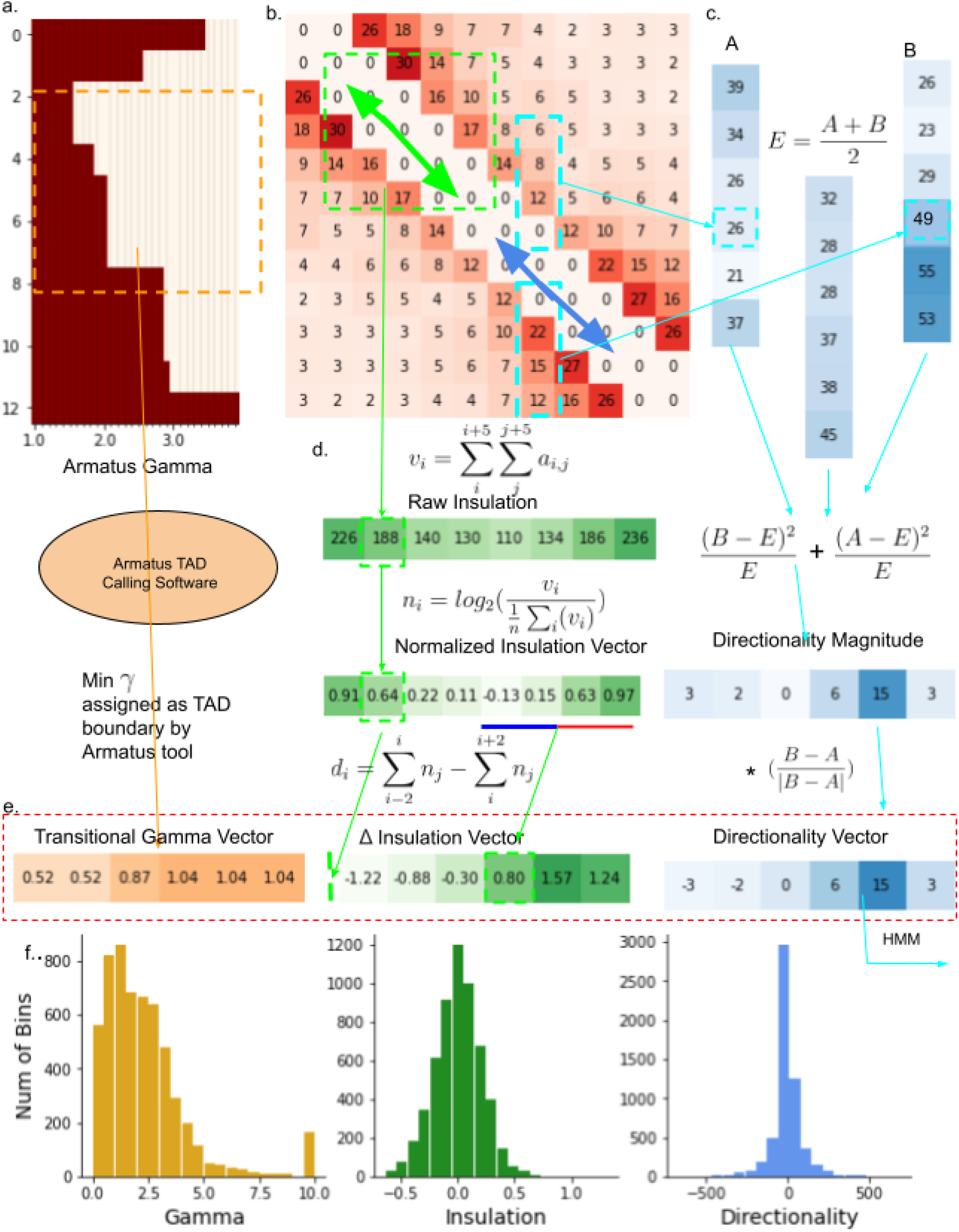
Dataset Overview. (a) Generation process of transitional gamma labels. (b) Hi-C Contact matrix (c) Generation process of Directionality Index labels. (d) Generation process of Insulation score labels. (e) vectors used as labels in the training process. (f) Distribution of numerical values for each metric

In addition to using the standard transitional gamma, we use Hi-C data (Figure 1b) to extract two additional labels for TAD characterization, which are prolific in the Hi-C literature: Directionality Index and Insulation Score.

We compute directionality index using a procedure based on Dixon et al (Dixon et al. 2012). Directionality Index is motivated by the observation that downstream portions of a domain are highly biased towards interactions with upstream bins. Directionality Index is computed using Equation 1, where A is a quantity of reads mapped from the observed bin to R bins downstream; B is a quantity of reads mapped from an observed bin to R bins upstream, and E is the mean of B and A. The result is a 1 dimensional vector with values corresponding to each genomic loci within R bins of the chromosome border. In the original Directionality Index literature the directionality vector is then passed to a hidden markov model, however, for ease of comparison we treat the directionality vector as our labels. Our directionality uses a radius of 10.

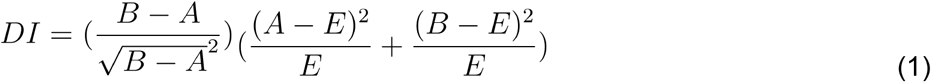

We compute the insulation score using a procedure based on (Crane et al. 2015). Insulation Score is motivated by the intuition that regions which have drastic changes in quantity of interactions with their neighbors are likely boundaries for TADS. Insulation Score (Figure 1d) is calculated by sliding a window of radius R along the diagonal of a contact matrix and computing the sum of signals across each bin. This vector is then normalized and a Difference Vector is computed by observing changes in the summed value of L bins before and after a loci of interest. The result is a 1 dimensional vector with values corresponding to each genomic loci within R+L bins of the chromosome border. In the original Insulation Score literature the regions of the Difference vector where values switch sign are marked as potential TAD boundaries. In our experiments we use the full vector as the label. We use R=3 and L=10.

### Overview of TAPIOCA Network

The TAPIOCA network is inspired by the transformer architecture originally proposed in 2017 as an approach to the problem of Seq2Seq language translation (Vazawani et al 2017) . To convert the transformer network from the task of language translation to TAD prediction, we treat epigenetic features as though they are word embeddings and append a final linear prediction layer converting transformer outputs to numerical values for TAD labels (Fig 2a). The core processing component of the TAPIOCA architecture is made up of a series of attention layers each containing Scaled Dot-Product Attention with 7 heads followed by a linear neural network layer (Fig 2b). Our network treats the number of attention layers as a hyper parameter which is tuned (Supplementary Figures 1-3).

**Figure 2.**
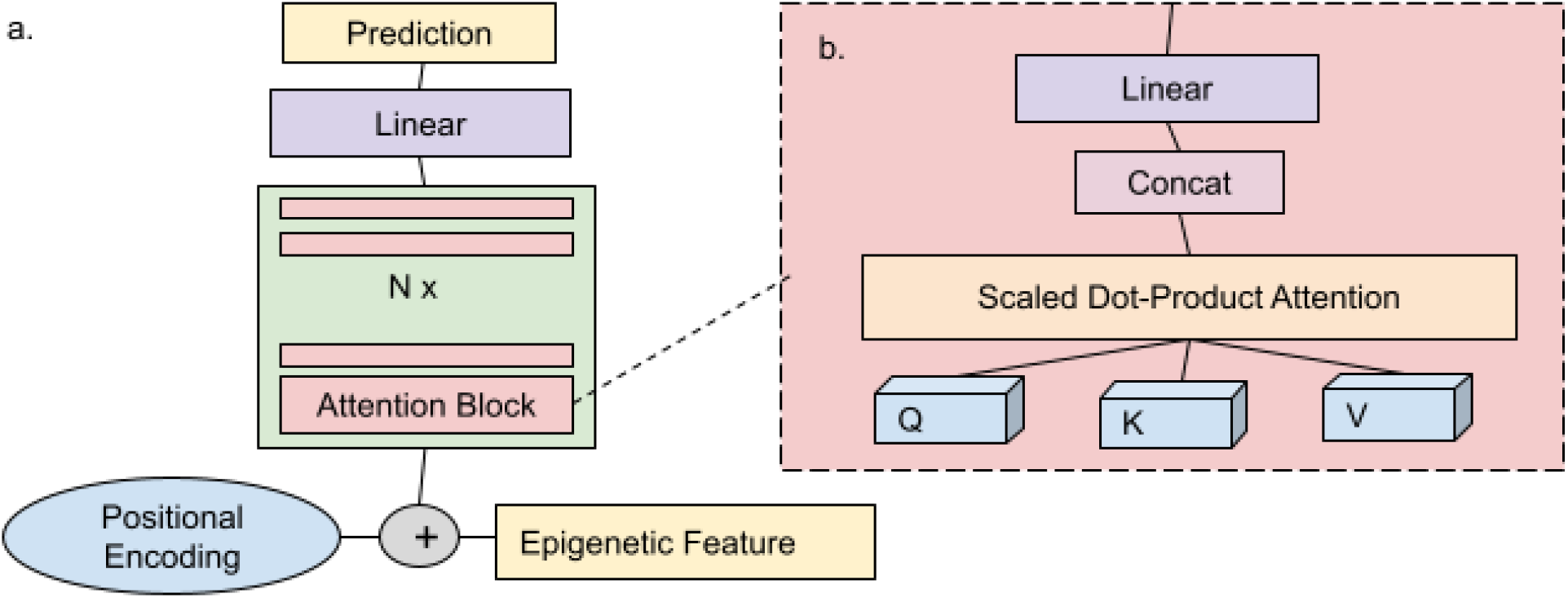
TAPIOCA Architecture. (a) Overview of architecture (b) Details of multi-head attention block.

**Figure 3.**
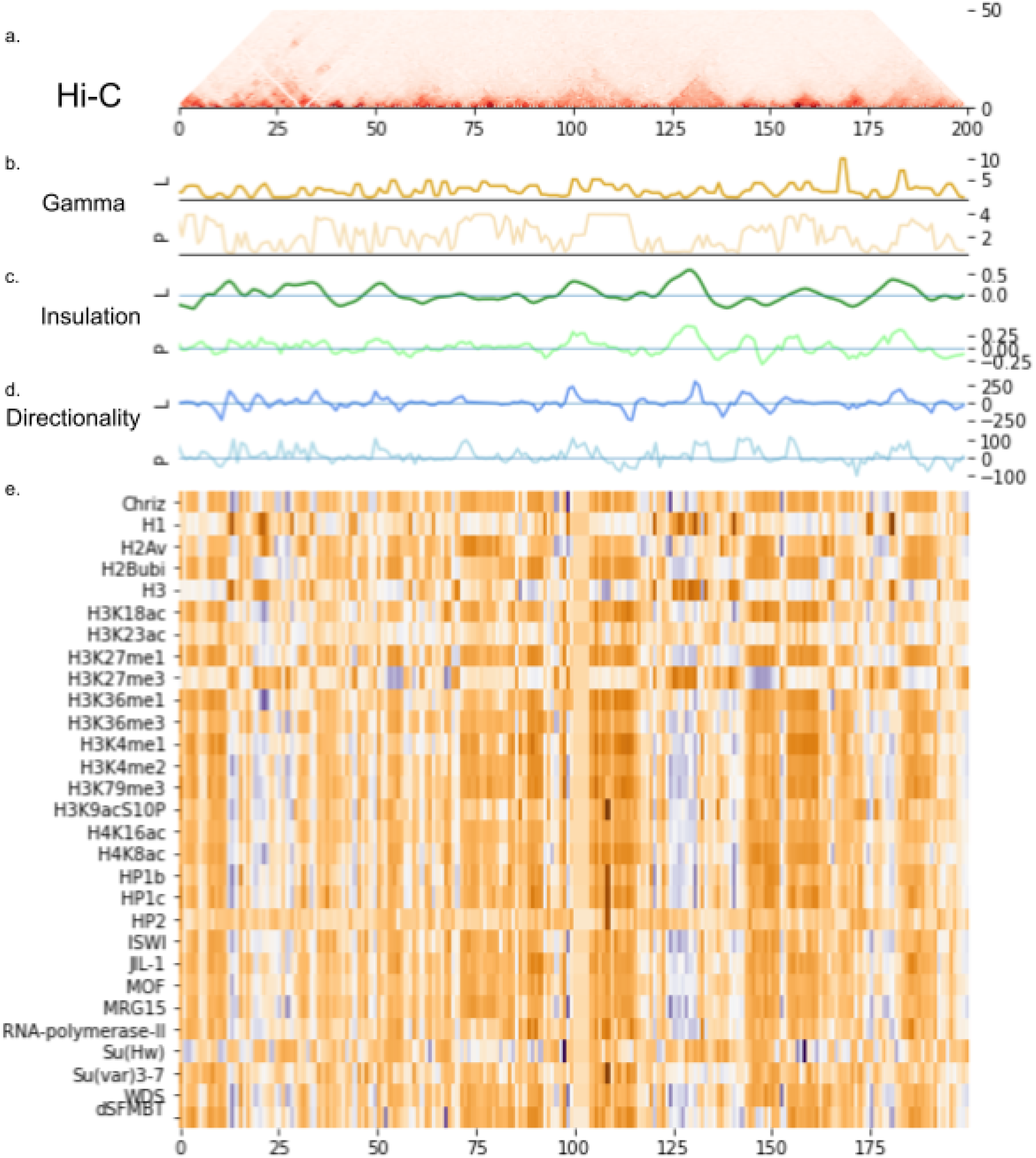
Visualization of Predictive Process. (a) Hi-C track data (b) Label and predicted values for transitional gamma, (c) insulation vector, and (d) directionality index. (e) epigenetic track data values.

### Benchmark of TAPIOCA Network Relative to Prior Art

We compared TAPIOCA-network’s performance on the task of TAD prediction to the performance of previously used models such as linear regression (Ulianov et al. 2016), ridge and lasso regression (Ramírez et al. 2018) and Bi-directional Long Short-Term Memory (BILSTM) (Rozenwald et al. 2020) We observe visual similarity between the predictions of our network and Hi-C derived labels across all three metrics (Fig 4). TAPIOCA-network outperforms all previous approaches on the transitional gamma dataset (Table 1). In insulation vector experiments TAPIOCA-network outperforms all linear regression variants while remaining competitive with BILSTM (Table 2). TAPIOCA-network was the only network capable of effectively predicting Directionality index, even after extensive hyperparameter tuning of other networks (Fig 4c, Table 3).

**Figure 4.**
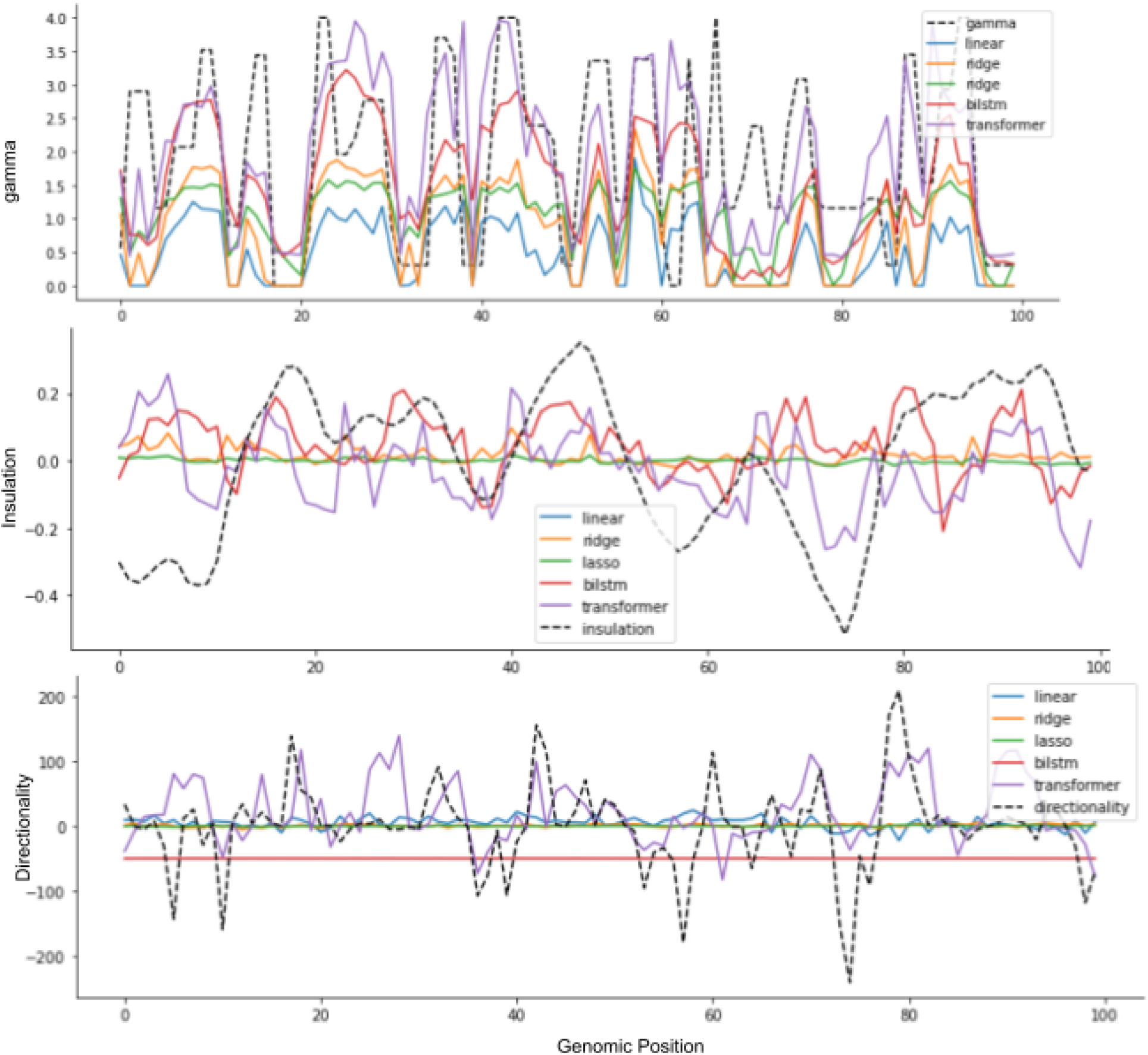
Benchmarking TAPIOCA. Predictions made by (blue) linear regression, (green) ridge regression, (yellow) lasso regression, (red) bidirectional long-short term memory network and (purple) tapioca network. The dotted black line shows Hi-C obtained predictive values for (a) transitional gamma, (b) insulation vector, and (c) directionality index.

**Table 1:**
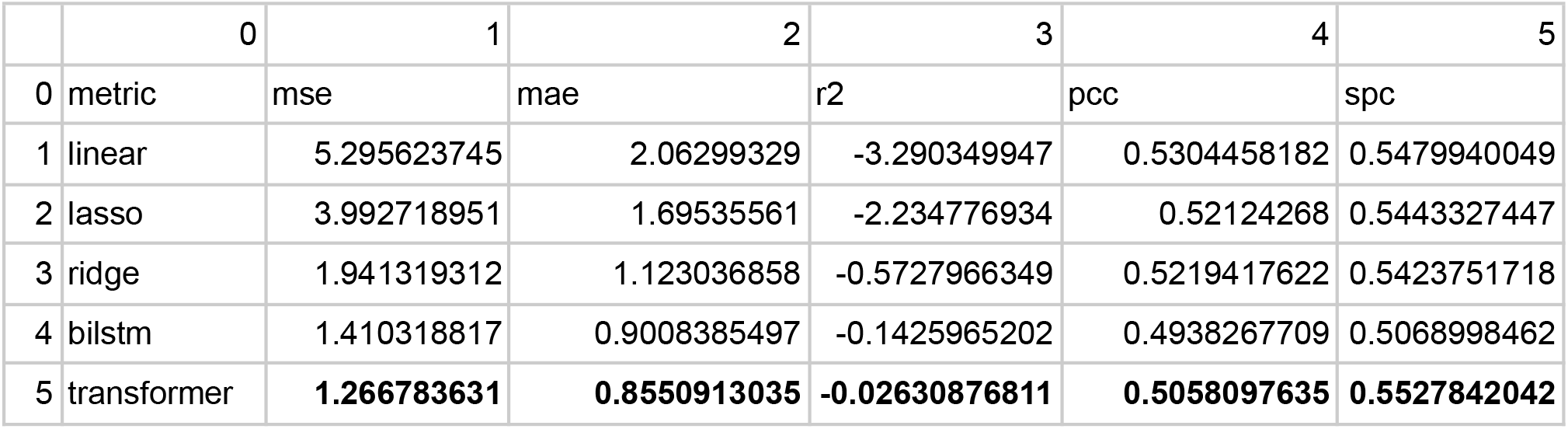
Performance metrics of different models on gamma using S2 cell line.

**Table 2:** Performance metrics of different models on insulation using S2 cell line.

|  | 0 | 1 | 2 | 3 | 4 | 5 |
| --- | --- | --- | --- | --- | --- | --- |
| 0 | metric | mse | mae | r2 | pcc | spc |
| 1 | linear | Na | NA | NA | NA | NA |
| 2 | lasso | 0.06974714665 | 0.2059695515 | -0.002145843473 | 0.058078392 | 0.07441855312 |
| 3 | ridge | 0.06951995737 | 0.204573979 | 0.00111847349 | 0.038189119 | 0.04004053408 |
| 4 | bilstm | **0.05829887199** | **0.1904818259** | **0.1623460593** | **0.419690222** | **0.3940576234** |
| 5 | transformer | 0.0652860434 | 0.2000407176 | 0.06195249305 | 0.324928991 | 0.3334798006 |

**Table 3:** Performance metrics of different models on directionality using S2 cell line.

|  | 0 | 1 | 2 | 3 | 4 | 5 |
| --- | --- | --- | --- | --- | --- | --- |
| 0 | metric | mse | mae | r2 | pcc | spc |
| 1 | linear | 8499.176481 | 57.25845387 | -0.01948713842 | -0.0392333315 | -0.02423517569 |
| 2 | lasso | 8349.548968 | 55.66149272 | -0.00153912602 | -0.0066974481 | 0.003306583346 |
| 3 | ridge | 8337.046141 | 55.49885533 | -3.94E-05 | 0.0185781542 | 0.01345918225 |
| 4 | bilstm | 10696.08042 | 78.29492895 | -0.2830085879 | -0.0551410883 | -0.02034854701 |
| 5 | transformer | **6534.42304** | **55.28723772** | **0.2161875611** | **0.5086738093** | **0.5152162301** |

### TAPIOCA Network Remains Effective Across Cell Lines

One of the most important characteristics of any machine learning algorithm is its ability to generalize. To ensure that our network’s predictive ability is not constrained to cell lines for which Hi-C data is already available, we test effectiveness of TAPIOCA at predicting TAD organization on cell lines which differ from the network training dataset. We observe that in most instances the network’s performance remains high even when training and test cell lines differ (Fig 5), in certain instances performing marginally better using different cell lines. Gamma obtains high values for R2 across all 5 metrics regardless of train test combination. The insulation vector and directionality index metric also obtain comparable results across training test cell line combinations in most instances.

**Figure 5.**
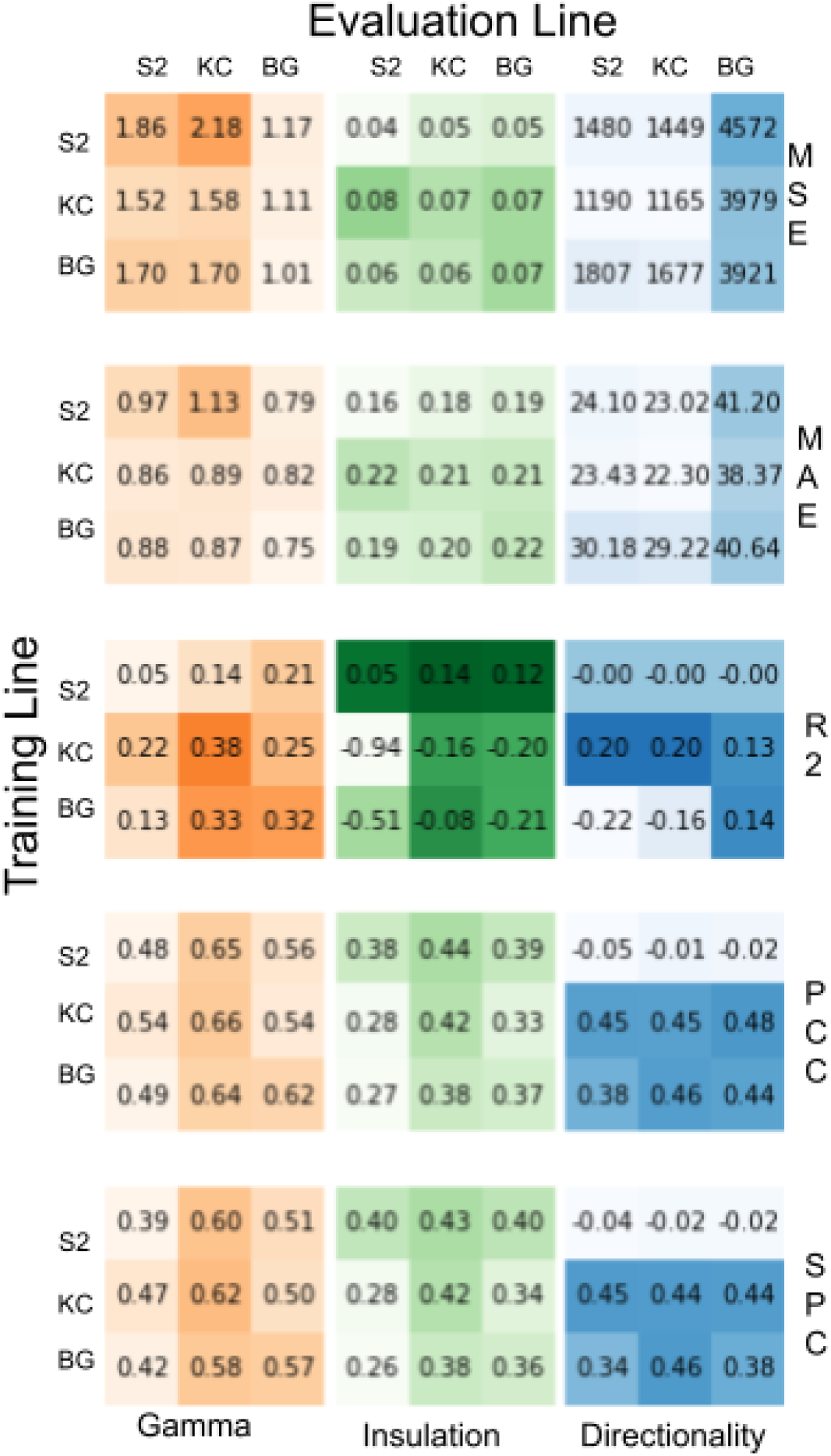
Performance Across Cell Lines. Rows indicate training set cell lines, columns indicate testing set cell lines using (orange) transitional gamma, (green) insulation vector, and (blue) directionality index labels. Super rows show metric of evaluation: mean squared error, mean average error, r2, pearson correlation and spearman correlation

### Key Epigenetic Features in TAD prediction using TAPIOCA Network Resembles the Key Features Observed in Prior Art

We ran experiments excluding each epigenetic feature from training in both our TAPIOCA network and the previous state of the art BiLSTM network. We observe high consistency in the evaluated performance of the TAPIOCA network across the metrics: mean average error, mean squared error, pearson correlation and spearman correlation (Fig 6a). We observe similar degradation of performance across both networks when excluding features (Fig 6b). Removing certain epigenetic features such as Chriz and Su(HW) showed sharp relative drops in performance on both networks, however, other previously identified features such as H3K27me3 and H3K27ac (Rozenwald et al. 2020) showed low relative degradation in performance of the TAPIOCA network.

**Figure 6.**
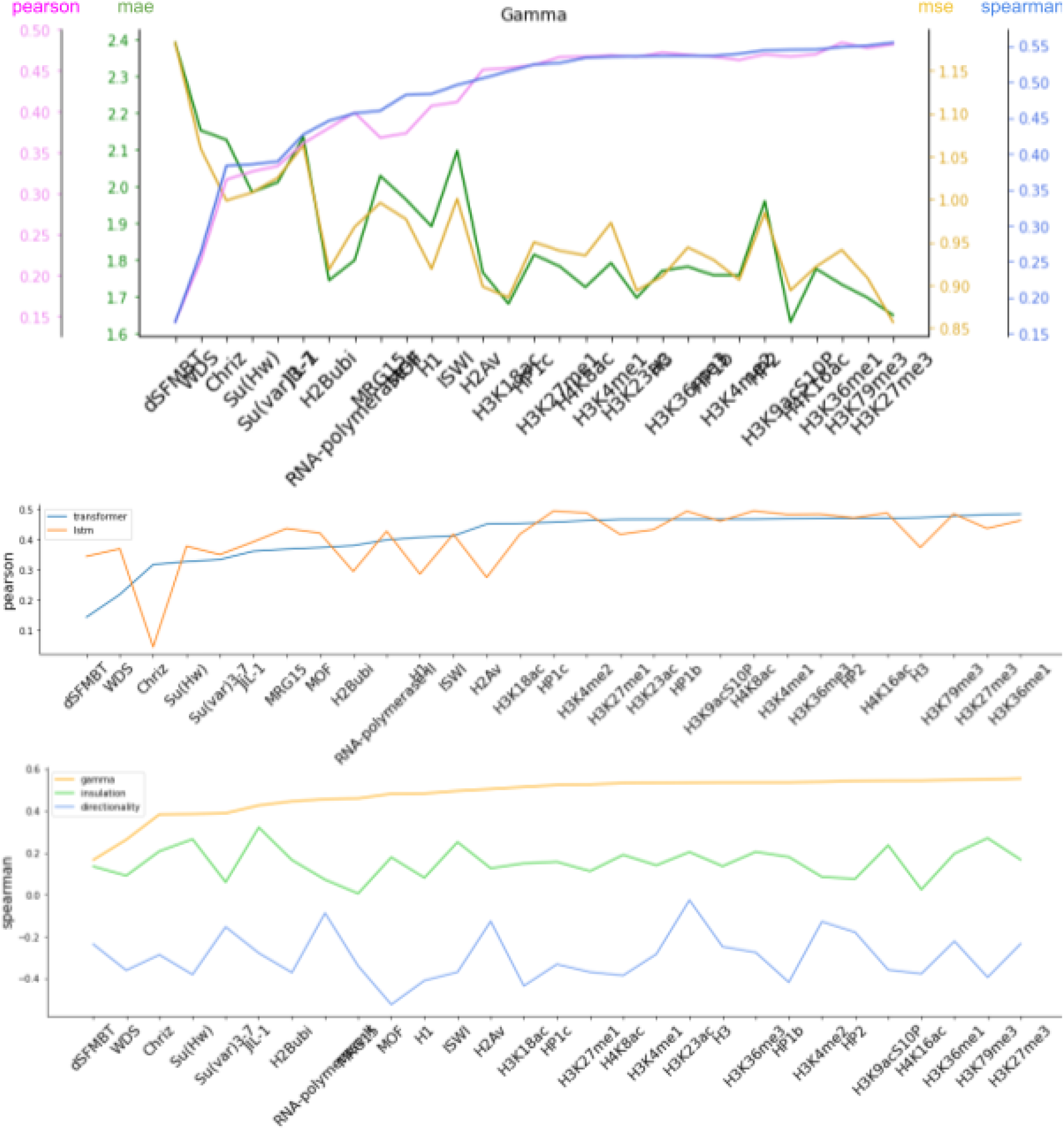
Removal of Features. (a) performance of TAPIOCA network when excluding single epigenetic features on (pink) pearson correlation, (green) mean average error, (yellow) mean squared error, and (blue) spearman correlation. (b) Pearson correlation of (blue) TAPIOCA network and (orange) bidirectional Long Short-Term memory network when excluding single epigenetic features . (c) Spearman correlation of TAPIOCA network predictions of (orange) transitional gamma (green) insulation score and (blue) directionality index when excluding single epigenetic features.

### Epigenetic Features have different priority in predictive ability based on TAD label selection

We ran experiments excluding each epigenetic feature across the three TAD identifying metrics (Fig 6c). In each case networks were trained using the hyperparameters which obtained best results in full features experiments. In many instances the directionality networks failed to converge to meaningful results. This may be due to the directionality datasets demonstrated difficulty and requirement for extreme hyper parameter tuning. While the insulation networks and gamma networks both converged in most feature exclusion experiments, the prioritization of values for epigenetic features showed little correlation (Fig 6c). While removing features such as dSFMBT, WDS and CHRIZ all showed pronounced decreases in performance for gamma prediction, the performance was not noticeably lower for these features in insulation score prediction relative to the removal of other epigenetic features such as H3K27Me3 and H3K27Ac.

## Discussion

We observe state of the art performance by TAPIOCA network on the well established metric of transitional gamma, indicating that the TAPIOCA approach should be considered when predicting TADs using epigenetic features. Furthermore TAPIOCA’s high performance on Insulation score and its unique success in Directionality index demonstrate the power of the transformer approach to modeling the complex predictive relationship of epigenetic profile and chromatin topology.

TAPIOCA’s ability to generalize across multiple cell lines indicates potential for real utility in saving the cost of Hi-C experiments, as this demonstrates that TAPIOCA can be used even without available Hi-C contact matrices from which to obtain labels. Future work could include examination of TAPIOCA’s effectiveness across other model organisms beyond Drosophila. Such experiments may provide insight to similarity of the underlying biological mechanism of TAD formation in different organisms.

In our experiments where we removed individual features, we observed a different set of epigenetic features whose absence maximally degraded model performance when using the TAPIOCA network than when using the previously described BILSTM. This seems to indicate that those decreases in performance may be unelucidated consequences of the selected machine learning models, rather than true biological relationships, raising uncertainty to the role of histone modifications H3K27Me3 and H3K27Ac in TAD formation as suggested in previous literature (Rozenwald et al. 2020). However, the epigenetic features where degradation of model performance was consistent between BILSTM and TAPIOCA, provide increased reason for hypothesizing an underlying relationship between TAD formation and presence of features such as Chriz.

We observed differing impacts on removing epigenetic features across different TAD characterization metrics. This disparity has multiple potential explanations and must be considered in conjunction with a few important observations. First, the explicit characterization of what makes a TAD is still an open area of discussion as multiple tools exist for TAD characterization and different tools often do not give fully concordant depictions (Zufferey et al. 2018). Second, the low performance of directionality index experiments in all epigenetic removal experiments is likely indicative of a failure of the model to capture the underlying data distribution, rather than anything grounded in biological reality. This claim is made in consideration with the demonstrated high hyperparameter sensitivity of the directionality dataset and the inability of BILSTM or regression variants to make successful predictions.

With these considerations, we do still observe differences in the contribution of removing single epigenetic features to successful prediction of insulation score and transitional gamma. One potential explanatory hypothesis for this disparity may be that different epigenetic features contribute to different scales or motifs of chromatin organization, some of which are more easily captured by armatus than insulation scores. Further work aiming to investigate this hypothesis may benefit from expanding analysis even further to include some of the many other TAD characterization metrics outlined in Zufferey et al. (Zufferey et al. 2018)

## Conclusion

In this manuscript we present, TAPIOCA, a tool for predicting TADs using epigenetic data via a self-attention based deep learning architecture. By reformulating the task of TAD prediction as a sequence transduction problem and developing an architecture inspired by the novel transformer network from machine learning literature we obtain state-of-the art results in inferring TAD characterization from epigenetic data. In addition to these results we contribute to the research community by expanding multiple Drosophila cell line datasets to include metrics for insulation score and directionality index.

## Methods

### Data

All data is based on cell lines from the Drosophila model organism. We use three cell lines: Schneider-2 (S2) and Kc167 from late embryos and DmBG3-c2 (BG3) from the central nervous system. Epigenetic profiles and transitional gamma labels for all cell lines were found at https://github.com/MichalRozenwald/Hi-ChIP-ML. Hi-C data used to construct Insulation and Gamma labels is available on the Gene Expression Omnibus at GSE69013. Cleaned datasets for all three metrics are available, along with all of our experiments at https://github.com/Max-Highsmith/TAPIOCA.

### Evaluation Metrics

We utilize 5 metrics for evaluation of similarity of predictions to labels: Mean Squared Error, Mean Absolute Error, R Squared, Pearson Correlation Coefficient and Spearman Correlation Coefficient.

#### Mean Squared Error

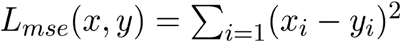

#### Mean Absolute Error

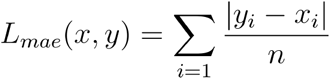

#### Coefficient of Determination

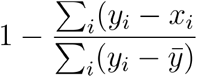

#### Pearson Correlation Coefficient

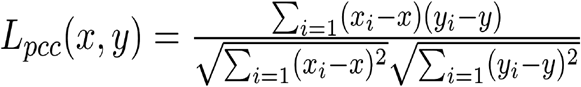

#### Spearman Correlation Coefficient

Spearman Correlation is similar to pearson correlation differing in that it utilizes rank variables so as to evaluate monotonic relationship between the matrices without imposing a linearity condition that may not exist in nature.

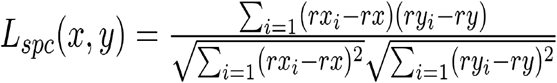

### Model Hyper Parameter Tuning

In all hyper parameter tuning experiments, we used a random search over sets of discrete values for each parameter. Hyper parameters were determined separately for each network with each TAD label. With each network we tested batch sizes (1,4,16,64), learning rates (1e5, 1e4, 1e3, 1e2, 1e1). With BILSTM and TAPIOCA we varied dropout (0,0.1, 0.2,0.3,0.5,0.7). And layer number (1,2,3,4,5,6). With TAPIOCA we varied the number of hidden units (512, 1024, 2048) and with BILSTM we varied bias existence for (True, False). All Models were initially trained with 10 iterations of random search for hyper parameters. Because none of these initial 10 results for the directionality dataset converged when using regression variants and BILSTM, we expanded the random search size to 20 but still did not obtain convergence on any network except TAPIOCA.

### Model Architecture and Training Details

The TAPIOCA model is inspired by the transformer architecture (Alammar n.d.). The Transformer was originally proposed for the task of seq2seq sentence translation and while our task is similar, there are a few key differences which inspired adjustments to the TAPIOCA architecture.

First, when working with seq2seq sentence translation, the fundamental unit of a sequence is a categorical token, typically a word. The preliminary step in translation tasks is the conversion of tokens into numerical vector representations via embedding. In the task of TAD prediction we begin with normalized epigenetic feature vectors for each 20kb region instead of tokens. This removes the need for inclusion of an embedding step because we already have vector representations.

Secondly, because multihead attention does not use recurrence or convolution it permits increased ability to identify relationships between spatially distant features. While this characteristic is clearly advantageous, it also necessitates manual inclusion of positional information into propagated vectors. In the original transformer architecture this task is performed by adding a positional encoding vector to embedded inputs. In the original transformer the positional encoding vector is more information dense for certain vector components. The assumption is that because the embedding layer is high dimensional (512) that the necessary information will be passed, and multiple components can permit meaningful feature integration of position and embedding. However, in the TAD identification task we eschew embedding completely, instead using epigenetic feature vectors. Thus additive positional encoding would have potential to overwrite or give implicit preference to components which already encode specific information. To prevent this we instead concatenate the positional econding vector to the epigenetic features.

Third, Unlike seq2seq translation there is no variation in sentence length of inputs and outputs. When training we use a sentence length of 11 bins, (220kb region). We use the mean squared error of our full predicted vector and label vectors as a loss function. When evaluating performance on test data we pass each sequence through with stride 1, keeping the middle vector bin.

## Acknowledgments

This project is supported by the NIH through the T32 training grant GM008396.

## Author Contributions

MH and JC conceived of the project. MH performed all experiments and drafted the manuscript. JC revised the manuscript

